# One to host them all: genomics of the diverse bacterial endosymbionts of the spider *Oedothorax gibbosus*

**DOI:** 10.1101/2022.05.31.494226

**Authors:** Tamara Halter, Stephan Köstlbacher, Thomas Rattei, Frederik Hendrickx, Alejandro Manzano-Marín, Matthias Horn

## Abstract

Bacterial endosymbionts of the groups *Wolbachia, Cardinium* and *Rickettsiaceae* are well-known for their diverse effects on their arthropod hosts, ranging from mutualistic relationships to reproductive phenotypes. Here, we analyzed a unique system in which the dwarf spider *Oedothorax gibbosus* is co-infected with up to five different endosymbionts affiliated with *Wolbachia, ‘Candidatus* Tisiphia’ (formerly Torix group *Rickettsia), Cardinium,* and *Rhabdochlamydia.* Using short-read genome sequencing data, we show that the endosymbionts are heterogeneously distributed among *O. gibbosus* populations and are frequently found co-infecting spider individuals. To study this intricate host-endosymbiont system on a genome resolved level, we used long-read sequencing to reconstruct closed genomes of the *Wolbachia, ‘Ca.* Tisiphia’ and *Cardinium* endosymbionts. We provide insights in the ecology and evolution of the endosymbionts and shed light on the interactions with their spider host. We detected high quantities of transposable elements in all endosymbiont genomes and provide evidence that ancestors of the *Cardinium, ‘Ca.* Tisiphia’ and *Wolbachia* endosymbionts have co-infected the same hosts in the past. Our findings contribute to broadening our knowledge about endosymbionts infecting one of the largest animal phyla on earth and show the usefulness of transposable elements as an evolutionary “contact-tracing” tool.

**Data summary:** All supporting data, code and protocols have been provided within the article or through supplementary data files. Seven supplementary figures and seven supplementary tables are available with the online version of this article. Sequencing data used in this study was generated and previously published by Hendrickx *et al.,* 2021. Genome assemblies generated in this study have been deposited under the project PRJEB52003 at DDBJ/ENA/GenBank. The MAG of *R. oedothoracis* OV001 was deposited at DDBJ/ENA/GenBank under the sample SAMN28026840. The genome of *‘Candidatus* Rhabdochlamydia oedothoracis W744×776’ was previously published by Halter *et al.,* 2022 and is available at DDBJ/ENA/GenBank (accession: CP075587-CP075588). The collection of genomes and proteomes, all files for phylogenetic analyses including gene alignments, concatenated alignments, and tree files, and original output files of the HGT and SNP predictions used in this study are available at zenodo (https://doi.org/10.5281/zenodo.6362846).

## Introduction

Arthropods are the largest and most diverse animal phylum on the planet [1], and they are frequently infected with endosymbiotic bacteria. Endosymbionts can have a variety of influences on their arthropod hosts including the manipulation of sexual reproduction, nutrition or resistance to pathogens [2]. Maternally transmitted endosymbionts belonging to the groups *Wolbachia, Cardinium,* and *Rickettsiaceae* are of particular interest because of the ability of some strains to induce reproductive phenotypes in their animal hosts [3–7]. These bacteria are also the most prevalent arthropod endosymbionts infecting around a half, a quarter and an eighth of terrestrial arthropod species, respectively [2].

The genus *Wolbachia* (Alphaproteobacteria) includes primarily maternally-transmitted endosymbionts of arthropods and nematodes, organized into at least 17 phylogenetic supergroups, where supergroup A and B are the most-well studied ones [3]. *Wolbachia* endosymbionts are most famous for the array of reproductive manipulations they can cause in their hosts, ranging from cytoplasmic incompatibility to male killing, parthenogenesis, and feminization [3]. However, they can also act as obligate or facultative mutualistic symbionts and have been reported to increase resistance against viruses in some host species [3]. Like *Wolbachia,* members of the genus *Cardinium* (Bacteroidetes) are endosymbionts of arthropods and nematodes [8]. Apart from causing reproductive manipulations [4, 5, 9, 10] *Cardinium* can be involved in the provision of nutrients [11] and in shaping its host’s microbiome [12, 13]. The family *Rickettsiaceae* (Alphaproteobacteria) includes the well-known arthropod-transmitted vertebrate and human pathogens of the genus *Rickettsia* but also invertebrate-restricted endosymbionts that are not transferred to vertebrate hosts and were recently placed in a separate genus *‘Ca.* Tisiphia’ (formerly Torix group *Rickettsia)* [14, 15]. Like *Wolbachia* and *Cardinium, ‘Ca.* Tisiphia’ include not only reproductive parasites that manipulate their host’s reproduction [6, 7, 16, 17] but also mutualists that can increase resistance to pathogenic fungi, bacteria and viruses [18–20].

Based on ribosomal RNA gene sequence analysis, the microbiome of the spider *Oedothorax gibbosus* was previously shown to be dominated by endosymbionts belonging to the above-mentioned genera and a less prevalent symbiont belonging to *Rhabdochlamydia* [21]. The genus *Rhabdochlamydia* belongs to the phylum Chlamydiae which consists of obligate intracellular bacteria [22] including major human and vertebrate pathogens, such as *Chlamydia trachomatis,* as well as symbionts of amoeba, and diverse environmental lineages with unknown hosts [22]. Known *Rhabdochlamydia* species are primarily endosymbionts of arthropods [21, 23–25], but their influence on the host remains to be investigated. However, there are indications that they have rather pathogenic than mutualistic effects [25–27]. Apart from harboring mainly endosymbionts, *O. gibbosus* was previously shown to be coinfected with multiple endosymbionts at the same time [21], which was only described for a few other hosts before.

Like for *O. gibbosus* most previous studies looking at spider endosymbionts have so far focused on describing the microbiome composition [21, 28–32]. However, whole genome information is needed to get deeper insights into host-symbiont interactions and to learn more about the ecology and evolution of the endosymbionts [33–36]. Here we reconstructed the complete genomes of four endosymbionts of the spider *O. gibbosus* including two different *Wolbachia,* one ‘*Ca.* Tisiphia’ and one *Cardinium* genome. Together with the previously published genome sequence of *‘Candidatus* Rhabdochlamydia oedothoracis’, we analyzed co-existence patterns of the endosymbionts across different *O. gibbosus* populations using short-read sequence data. We then compared the endosymbiont genomes to their closest known relatives and studied nutrient provisioning pathways, secretion systems, and potential effectors of the endosymbionts to better understand host-microbe and microbe-microbe interactions in this system. The complete and closed symbiont genomes also enabled us to have a closer look at transposable elements. By reconstructing their phylogenies, we provide a glimpse into the evolutionary past and found evidence for an early co-existence of the endosymbionts in the same hosts.

## Results and Discussion

### Co-occurence of endosymbionts in different host populations

Co-infections of multiple reproductive symbionts of the groups *Wolbachia, Cardinium* and *Rickettsiaceae* within the same individual appear to happen only rarely. To get an idea about the distribution of the endosymbionts among *O. gibbosus* individuals from geographically distinct populations, we first reconstructed the endosymbiont genomes from sequence data obtained from the host *O. gibbosus* by Hendrickx *et al.,* 2021 (Table S1). We calculated the relative abundance of the symbionts and symbiont load (*i.e.* number of symbionts per diploid host cell) by mapping short-read sequence data obtained from adult male *O. gibbosus* individuals to the reference genomes of the endosymbionts.

We observe co-infections with multiple endosymbionts to occur frequently in *O. gibbosus* as 10 out of 16 spiders from six different locations across Belgium are infected with two or more endosymbionts. Further, we noticed a large variability regarding the distribution of the endosymbionts even between *O. gibbosus* individuals of the same population (Figure 1). *Cardinium* is the most prevalent symbiont occurring in almost all sequenced individuals (94%), while *Rhabdochlamydia* is limited to the Walenbos and Overmeren population (Figure 1). Further, except for individuals containing *Rhabdochlamydia,* the symbiont load is generally low (3.10 symbionts per host cell on average) (Figure 1). In spiders infected with *Rhabdochlamydia* the number of endosymbionts increases to 42.5 on average, with *Rhabdochlamydia* being the most abundant member of the community. Its high abundance could point towards a pathogenic rather than mutualistic or commensal role of *Rhabdochlamydia* in the spiders as suggested for other members of the genus [25–27]. In addition, we found the two *Wolbachia* symbionts co-infecting individuals in two different populations. Interestingly, we found *Wolbachia* in two male spiders from the Walenbos population (Figure 1) that was previously shown to be affected by male-killing *Wolbachia* [37]. As suggested earlier, this could hint towards an incomplete male-killing allowing some male offspring to survive [38].

**Figure 1:**
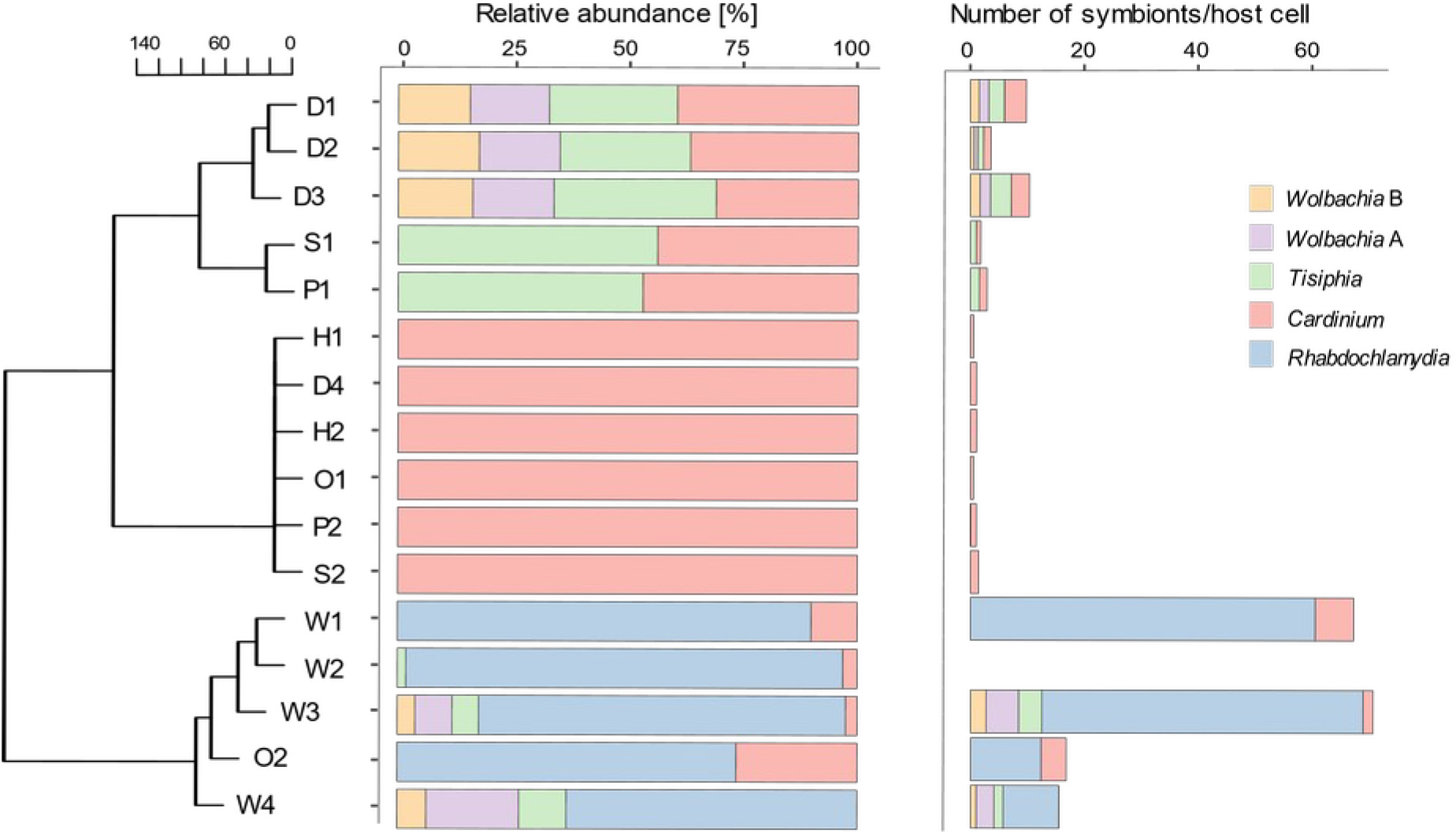
Relative abundance of the endosymbionts and symbiont load in male *O. gibbosus* individuals of different populations. Relative abundances and symbiont load *(i.e.,* number of symbionts per diploid host cell) were calculated based on the coverage of the endosymbionts normalized by the coverage of elongation factor alpha of *O. gibbosus.* The samples were grouped using hierarchical clustering based on an euclidean distance matrix. Samples were obtained from different locations across Belgium: D=Damvallei (n=4), S=Sevendonck (n=2), P=Pollismolen (n=2), H=Honegem (n=2), O= Overmeren (n=2), W=Walenbos (n=4).

To get an idea about the genetic variability between the endosymbionts of different populations we analyzed genomic variations, i.e. single nucleotide polymorphisms (SNP), insertions/deletions (indels), and substitutions by mapping short read sequence data from individual spiders against the reference genomes obtained from the Walenbos population. Whereas we find only low numbers of variants (0.1 mutations/kB on average) for *Cardinium* and *Rhabdochlamydia* suggesting nearly clonal symbiont populations, the ‘*Ca.* Tisiphia’ symbionts are considerably more heterogeneous (7.1 mutations/kB on average) (Figure S1).

### Genomic repertoire of the endosymbionts

#### Rhabdochlamydia

The genome of the *Rhabdochlamydia* symbiont of *O. gibbosus* was previously described by Halter *et al.,* 2022. It revealed a high similarity to its closest relatives *R. porcellionis* and ‘*Ca.* R. helvetica’, infecting crustacean and arachnid hosts, respectively [23, 24]. Unlike other *Rhabdochlamydia*, the genome of ‘*Ca.* R. oedothoracis’ contains a large number of transposable elements making up about 23% of the genome and leading to the hypothesis that its genome is currently undergoing genome size reduction [39]. Here, we compare the published genome of ‘*Ca.* R. oedothoracis W744×776’ (hereafter *R. oedothoracis)* isolated from *O. gibbosus* from the Walenbos population with a medium-quality metagenome assembled genome (MAG, sensu [40]) (Table S1) newly reconstructed from *O. gibbosus* from the Overmeren population (‘*Ca.* R. oedothoracis OV001’, hereafter MAG rhOegib-Ov). According to the average amino acid identity (AAI; 99.47 %) and average nucleotide identity (ANI; 99.79 %) the two genomes belong to the same species [41].

To compare the two genomes we first clustered all predicted proteins into orthologous groups (OG; corresponding to gene families) using either eggNOG (v4.5) [42] or silix (v1.2.11) [43], where silix was used for *de-novo* clustering of genes not annotated in eggNOG (v4.5) [42]. For the best studied members of the phylum Chlamydiae, namely members of the family Chlamydiaceae, it has previously been shown that strains of the same species are highly similar in gene content and genome synteny [44–46]. We could observe a similar pattern for *R. oedothoracis*, as the two genomes share most of their gene families (Figure 2A). The number of accessory gene families that are specific for one of the genomes is significantly larger for *R. oedothoracis.* However, we observed that most of the accessory genes of *R. oedothoracis* are located in close proximity to transposase genes (Figure 2B), suggesting that those genes were probably not assembled in MAG rhOegib-Ov due to the presence of multiple copies of diverse transposases. Such assembly challenges in metagenomic data highlights the importance of available complete genome sequences especially for groups known to include many transposable elements. Our comparative analysis further confirmed the involvement of transposable elements in genome rearrangements and genome evolution in *Rhabdochlamydia oedothoracis* as suggested earlier [39], as the location of transposase genes tend to correspond to breakages in synteny between the two genomes (Figure 2C).

**Figure 2:**
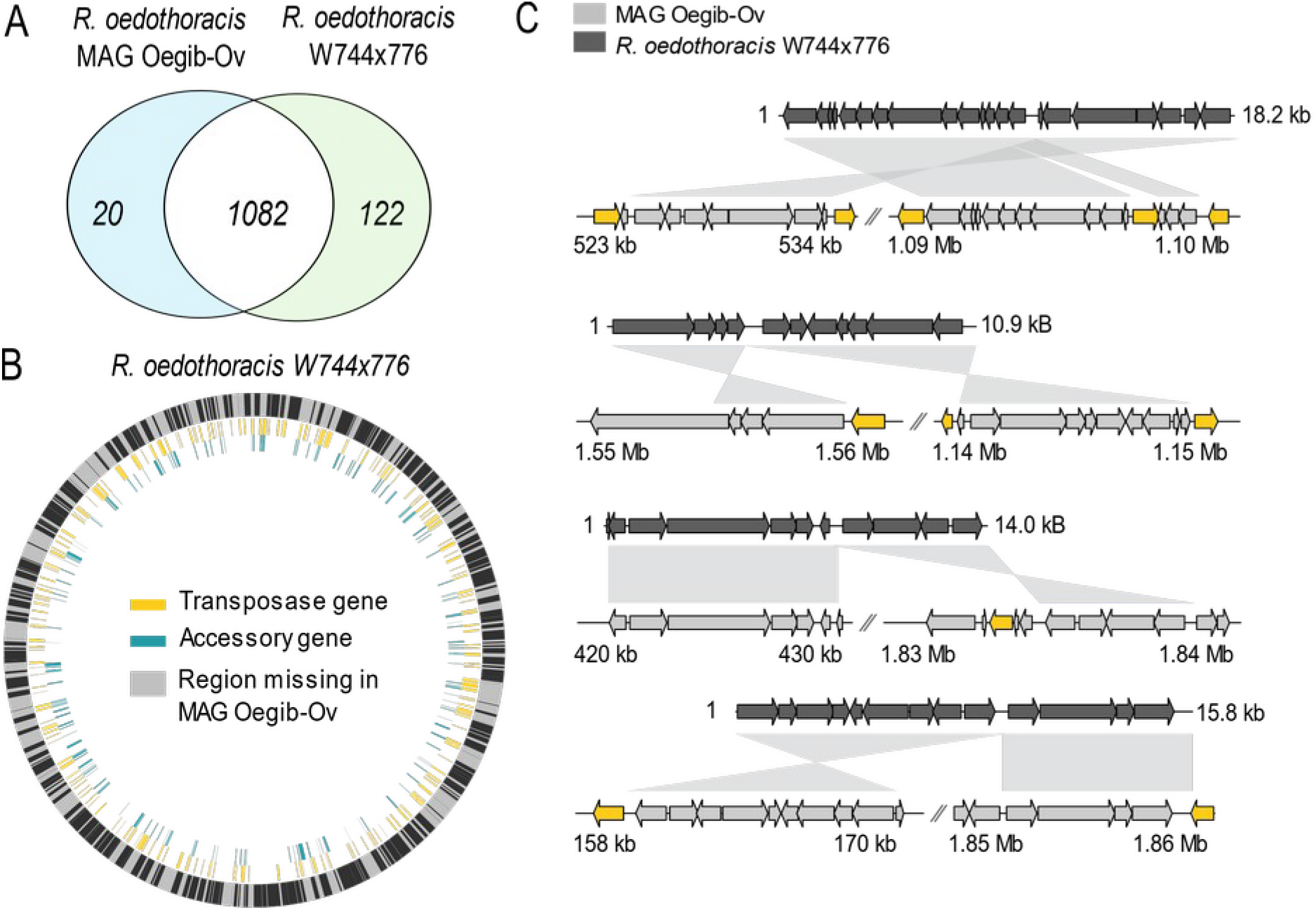
Genome comparison of two *Rhabdochlamydia oedothoracis* strains, W744×776 and MAG rhOegib-Ov. (A) Genome content comparison based on eggNOG and *de-novo* clustered gene families (OGs). For the analysis only proteins with a minimum length of 100 amino acids were used. (B) Genome alignment of *R. oedothoracis* W744×776 and MAG rhOegib-Ov. Regions present in MAG rhOegib-Ov are shown in black, annotated transposases in the genome of *R. oedothoracis* W744×776 are shown in yellow, and accessory genes of *R. oedothoracis* W744×776 are shown in cyan. (C) Selected regions representing genomic rearrangements and co-localized transposase genes between the *R. oedothoracis* W744×776 and MAG Oegib-Ov. Transposase genes are shown in yellow and other CDSs in shades of grey.

#### Cardinium

The genome of the endosymbiont *Cardinium* sp. cOegibbosus-W744×776 (hereafter cOegib-Wal) (Table S1) is 1.1 Mb in size and contains a high number of transposable elements, making up around 30% of the genome. The ribosomal protein phylogeny places *Cardinium* sp. cOegib-Wal in the group A *Cardinium* (Figure 3A) [47, 48], which encompasses *Cardinium* strains associated with insects and arachnids [47]. *Cardinium* sp. cOegib-Wal is nested in a clade with *Cardinium* symbionts of the rice bug *Stenocoris furcifera* (cSfur) and the mite *Dermatophagoides farinae* (cDfar) that seems to consist of three distinct species based on both ANI and AAI (Table S2). The sister clade comprises *Cardinium* symbionts of the whitefly *Bemisia tabaci* (cBtQ) and the parasitic wasp *Encarsia pergandiella* (cEper). Their genomes are very similar on nucleotide level and based on their genome features even suggested to be strains of the same species [49]. However, this finding is not surprising as there is an ecological link between *Cardinium* sp. cBtQ and *Cardinium* sp. cEper: *E. pergandiella* is a common parasitoid of *B. tabaci*, while the host of the other *Cardinium* group A members are not ecologically related. The closest sequenced relative to *Cardinium* sp. cOegib-Wal is the widely distributed mite-associated *Cardinium* sp. cDfar [50] (Figure 3A). In mites, *Cardinium* may lead to a sex-ratio bias as it was previously found more abundant in females than in males (67% and 92%, respectively) [50]. However, little is known about sexratio bias in astigmatid mite species, and to our knowledge there are also no candidate genes for *Cardinium* associated with reproductive manipulation of these hosts. However, it is important to note, that previous studies in *O. gibbosus* did not find a correlation between the presence of *Cardinium* and a sex-ratio bias [37].

**Figure 3:**
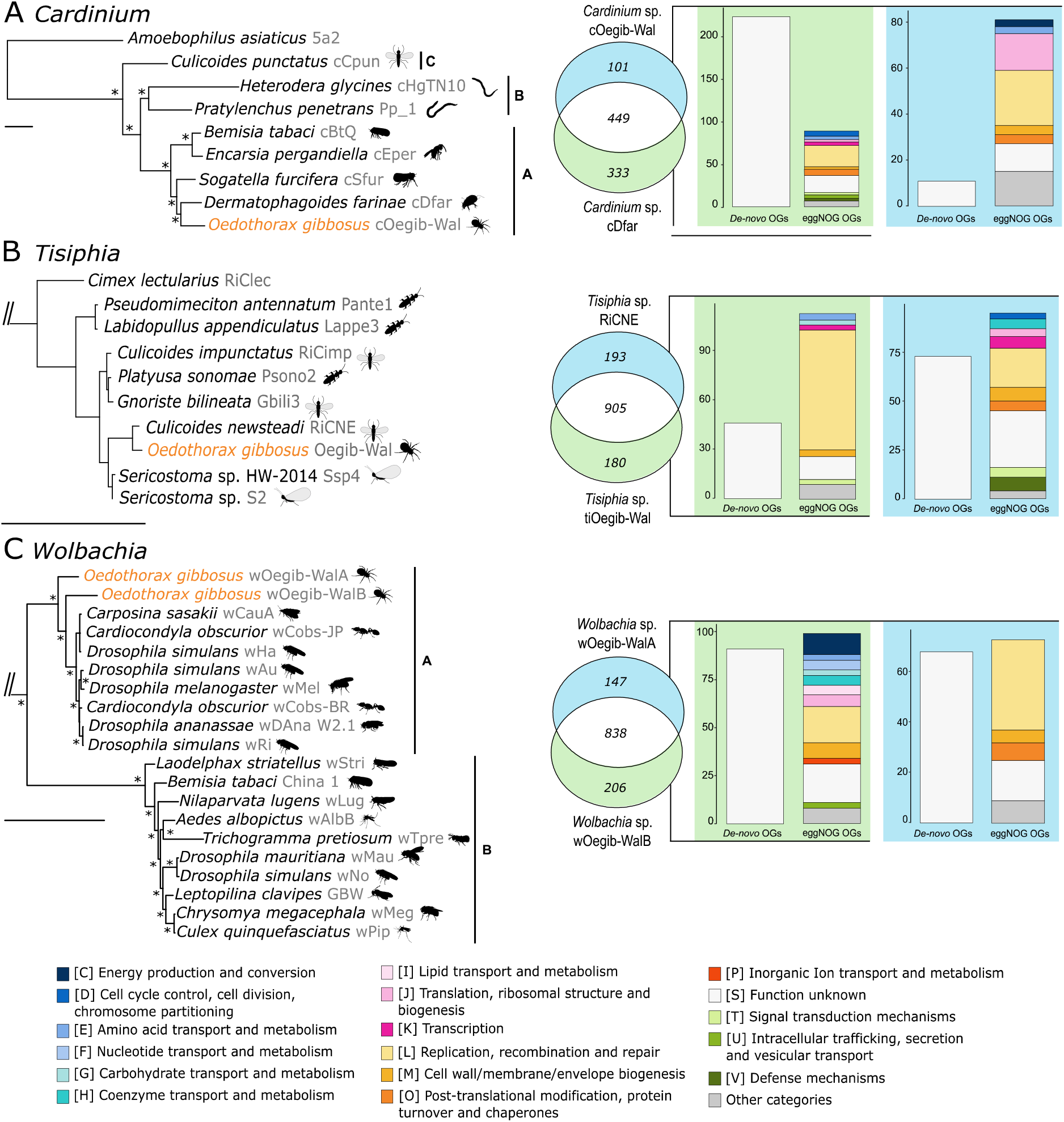
Relationship of the *Cardinium, ‘Candidatus* Tisiphia’ and *Wolbachia* endosymbionts of *O. gibbosus,* and genome comparison with their closest relatives. (A) Ribosomal protein gene phylogeny of the genus *Cardinium.* Scale bar indicates 0.07 substitutions per position in the alignment. Asterisks mark branches with a posterior probability of 1. The tree was rooted using the amoeba symbiont *Amoebophilus asi*aticus 5a2 as an outgroup. (B) Ribosomal protein gene phylogeny of the genus *‘Candidatus* Tisiphia’. Scale bar indicates 0.1 substitutions per position in the alignment. The tree was rooted using *Orientia* sp. and *‘Candidates* Megaira’ sp. as an outgroup. All branches have a support value ≥95. (C) Ribosomal protein gene phylogeny of the genus *Wolbachia.* Scale bar indicates 0.07 substitutions per position in the alignment. Asterisks mark branches with a posterior probability of 1. The tree was rooted using *Anaplasma phagocytophilum* and *Ehrlichia canis* as an outgroup. Host names are indicated at the leaves. For clear presentation only selected clades are shown in the trees, the complete dataset used to construct the trees is shown in table S7. Genome comparison as Venn diagrams and composition of accessory genomes based on eggNOG and *de-novo* clustered gene families (OGs) are shown next to the phylogenetic trees. Only unambiguously annotated COG categories were used, and only categories n>2 are shown, the rest is clustered into “Other categories”. For the analysis only proteins with a minimum length of 100 amino acids were used.

Our genome content analysis of *Cardinium* sp. cOegib-Wal and its closest relative *Cardinium* sp. cDfar revealed a reduced genomic repertoire of cOegib-Wal and a substantial fraction of gene families that is specific for the mite symbiont cDfar (Figure 3A). The putative function of these gene families was inferred from eggNOG (v4.5) [42] and blastp [51] against the NCBI-nr database. Most represent hypothetical proteins (COG category S and *de-novo* OGs) with homologs in other *Cardinium* strains, suggesting a loss of those genes in *Cardinium* sp. cOegib-Wal. The presence of COG categories involved in metabolic processes (C, D, E, J, K, O) in the accessory genomes of cOegib-Wal and cDfar likely indicate differences in the basic metabolism while the presence of COG categories involved in defense mechanisms, secretion, signal transduction and cell wall/membrane biosynthesis (V, U, T, and M) likely indicate adaptation to different hosts and environments (Figure 3A).

Notably, the genome of cOegib-Wal encodes a large non-ribosomal peptide synthase (NRPS) cluster consisting of a large modular protein and its helper proteins. NRPSs are widespread in prokaryotes and eukaryotes and of particular interest due to their antimicrobial and antifungal properties [52]. The NRPS found in cOegib-Wal is predicted to synthesize a short “TNTXXLP” peptide (Figure S2), for which its action remains unknown. From currently fully sequenced *Cardinium* genomes, only the nematode-infecting strain cHgTN10 contains a NRPS gene cluster. However, these two NRPSs are not homologous. In our analysis, *Cardinium* appears to be the most prevalent endosymbiont in *O. gibbosus* (Figure 1), suggesting a potential role of the NRPS in shaping the microbiome of its spider host.

#### *Candidatus* Tisiphia

The closed genome of *‘Candidatus* Tisiphia’ sp. Oegib-W744×776 (hereafter *Tisiphia* sp. tiOegib-Wal) is 2.6 Mb in size (Table S1) and consists of 28% transposable elements. According to the ribosomal protein phylogeny the genome belongs to the recently proposed genus ‘*Ca.* Tisiphia’ (formerly Torix group *Rickettsia*), a sister group of the genus *Rickettsia* [15] (Figure 3B). The phylogenetic placement is further corroborated by a 16S rRNA phylogeny (Figure S3). Its closest sequenced relative is ‘*Ca.* Tisiphia’ sp. RiCNE (formerly *Rickettsia* sp. RiCNE, hereafter *Tisiphia* sp. RiCNE), a symbiont of the midge *Culicoides newsteadi* (Figure 3B) [53]. According to the AAI (94.11 %) and ANI (97.00 %) scores, the two genomes belong to the same species. We could not observe a pattern regarding phylogeny and host range in ‘*Ca.* Tisiphia’ (Figure 3B, Figure S3). Further, *Tisiphia* sp. RiCNE and *Tisiphia* sp. tiOegib-Wal are strains of the same species but were isolated from phylogenetically distinct hosts. Together, this could indicate a propensity for horizontal transmission of this ‘*Ca.* Tisiphia’ species between host species.

Although the two strains share most of their gene families, about 17% of the gene families are unique to either one of them (Figure 3B). Most of the gene families missing in *Tisiphia* sp. tiOegib-Wal are hypothetical proteins with close homologs in other *Rickettsiaceae* suggesting a loss of those genes in *Tisiphia* sp. tiOegib-Wal. Furthermore, *Tisiphia* sp. tiOegib-Wal seems to have a more reduced genomic repertoire with its accessory gene families falling into only a few different functional categories (Figure 3B) and most of them being transposases and genes for their maintenance (COG category L). As shown for *R. oedothoracis* it is likely that those gene families are not assembled in the available draft genome of *Tisiphia* sp. RiCNE. Interestingly, only a few accessory gene families are linked to host interaction (Category T, M) (Figure 3B) although the two strains were isolated from very different hosts.

#### Wolbachia

We were able to reconstruct two different closed *Wolbachia* genomes (hereafter *Wolbachia sp.* wOeGib-WalA and *Wolbachia sp.* wOeGib-WalB) from *O. gibbosus.* According to the ribosomal protein gene phylogeny (Figure 3C), the AAI and ANI (89 % and 93 %, respectively; Figure S4, Figure S5, Table S4), the two genomes represent separate lineages falling at the base of supergroup A of *Wolbachia*. Compared to the AAI/ANI of other supergroup A and B *Wolbachia* they still belong to supergroup A and do not form new supergroups (Figure S4, Figure S5, Table S4). Both genomes show a substantial fraction of transposable elements (13% wOeGib-WalA, 12% wOeGib-WalB) consistent with reports about other *Wolbachia* genomes [54, 55].

Like previously shown for other *Wolbachia* lineages, *Wolbachia sp.* wOeGib-WalA and *Wolbachia sp.* wOeGib-WalB share most of their gene content (Figure 3C) [56]. Most accessory gene families in both genomes are hypothetical proteins (Figure 3C; *de-novo* OGs, COG category S) with close homologs in other *Wolbachia* suggesting a differential loss in one of the genomes. *Wolbachia sp.* wOeGib-WalB seems to have a larger and more diverse genetic repertoire as there are different functional categories represented in its accessory genome (Figure 3C). In *Wolbachia sp.* wOeGib-WalA, on the other hand, most gene families with an assigned function fall into COG category L, which includes transposases and genes for their maintenance.

### Interactions of the endosymbionts with the host

#### Reproductive manipulation

Although members of all three groups: *Cardinium*, *Wolbachia* and *Rickettsiaceae* were shown to be able to manipulate their hosts’ reproduction [3–7], a previous study only found a correlation between the presence of *Wolbachia* and female-biased offspring in *O. gibbosus* [37]. As the number of male and female eggs was similar in sex-biased and unbiased breeds, male-killing was suggested to cause the sex ratio bias [37]. *Wolbachia*-induce*d* sex ratio bias is thought to be mainly caused by the products of the *wmk* and the *cifA* and *cifB* genes [57]. Homologs of the *wmk* gene are present in all known male-killing *Wolbachia* strains [57], and there is growing evidence that this gene has a crucial role in male-killing [58]. We identified *wmk*-like genes in both *Wolbachia* genomes from *O. gibbosus.* These genes are most similar to the homologs in the *Wolbachia* sp. wInn symbiont of *Drosophila innubila* (wOegib-WalA) and *Wolbachia* sp. wBif infecting the moth *Drepanogynis bifasciata* (wOegib-WalB). However, recent evidence has shown that transgenic expression of the wInn or wBif *wmk* homologues in *Drosophila melanogaster* does not recapitulate the male-killing phenotype [58]. Nonetheless, given that the genes were expressed without the presence of the corresponding *Wolbachia* and in a non-native insect host, the male-killing phenotype of these *wmk* homologues cannot be ruled out in their native hosts. The *cifA* and *cifB* genes known to be involved in cytoplasmic incompatibility are absent in *Wolbachia* sp. wOegib-WalA, but we were able to identify three clusters of highly similar, and likely paralogous, *cifA* and *cifB*-like genes in *Wolbachia* sp. wOegib-WalB (Figure S6). However, these genes are quite divergent to *cifA* and *cifB* genes of other *Wolbachia* strains (average sequence identity *cifA* 19% and *cifB* 17%). Therefore, their actual role as cytoplasmic incompatibility factors remains to be tested. Genes associated with reproductive phenotypes i.e. *wmk*, *cifA* and *cifB* are often found in genomic islands that are associated with WO, a temperate bacteriophage that infects many *Wolbachia* strains [59, 60]. Interestingly, the genomes of *Wolbachia* sp. wOegib-WalA and *Wolbachia* sp. wOegib-WalB do not contain such an island and only exhibit traces of the WO phage.

#### Amino acid and vitamin synthesis

All five endosymbionts of *O. gibbosus* have reduced metabolic capabilities (Table S3). None of them is capable of synthesizing essential amino acids, and except for *Wolbachia* none of them is able to produce B-vitamins (thiamin, riboflavin, pyridoxal-5-phosphate, coenzyme A, folic acids, biotin). This observation is consistent with *O. gibbosus* being a generalist predator of small insects and thus not being limited in nutrients by its diet. Interestingly, *Wolbachia* sp. wOegib-WalA and wOegib-WalB encode for a complete biotin synthesis pathway (Table S3) which includes six genes *(bioC, bioH, bioF, bioA, bioD* and *bioB)* that are organized in a gene cluster. The genes show high sequence similarity to a complete biotin operon that was found in only five other *Wolbachia* strains yet and suggested to have been recently horizontally acquired by *Wolbachia* [61–63]. While the biotin operon is used by some *Wolbachia* strains to supplement their hosts’ vitamin B deficient diet [69, 71], the diverse diet of *O. gibbosus* is generally not regarded to be limited in biotin. However, as there is no information about the biotin needs of the spider, the role of biotin production by *Wolbachia* for the system remains to be investigated.

#### Toxin-antitoxin systems

Toxin-antitoxin (TA) systems are two-component systems composed of a stable toxin (protein) and its labile antitoxin (RNA or protein) [64], where most toxins interfere with protein biosynthesis [65]. Originally, TA systems were discovered as plasmid maintenance tools, killing daughter cells that lack a functional copy of the plasmid [66]. However, later on TA systems were identified on almost all bacterial chromosomes [66]. Although the function of these chromosomal copies is still unclear, it is hypothesized that they might regulate bacterial growth or operate in host cell manipulation [67].

The genome of *Tisiphia* sp. tiOegib-Wal contains three type-II TA modules: two DinJ/YafQ and one HigB/A. In type-II TA systems both the toxin and the antitoxin are small proteins that form a protein-protein complex resulting in the neutralization of the toxin. Further, there is one TA module with only low sequence similarity to any described TA system in other *Rickettsiaceae,* and four type-II toxins (YoeB, RatA, BrnT) and two antitoxins not arranged in modules (Table S4). Generally, it is unusual to find toxins without the respective antitoxin as it was previously shown that the overexpression of type-II toxins results in cell death in the absence of its antitoxin [67]. However, there are other described genomes only encoding toxins suggesting the presence of unknown mechanisms underlying antitoxin activity and a more complex relationship between toxin and antitoxin activities at physiological expression levels [67]. For *Tisiphia* sp. tiOegib-Wal we could identify potential antitoxins for the YoeB toxins encoded at the 5’ ends of the toxins. We found a similar module also in the genome of *Rickettsia felis,* and the putative antitoxin has close homologs in other *Rickettsia* and *Orientia.* However, there is no homology to the described antitoxin of YoeB, YefM.

The genome of *Wolbachia* sp. wOegib-WalB contains three copies of toxin RelE (Table S5) which is part of the RelEB type-II TA system also found in other *Wolbachia* genomes [68]. The three RelE copies are not orthologous according to eggNOG but contain a RelE domain. Nonetheless, we found no evidence for the presence of the corresponding antitoxin RelB that is present in other *Wolbachia* genomes.

#### Secretion systems and potential effectors

Secretion systems are used by bacteria to export substrates out of the bacterial cell. These so-called effectors serve to communicate with the environment and in case of intracellular bacteria with their host cells [69]. For the endosymbionts of *O. gibbosus* we could identify different secretion systems that are well described for the respective clade. *R. oedothoracis* encodes, like all known members of the phylum Chlamydiae, a complete type III secretion system [22, 39, 70]. The genome of *Cardinium* sp. cOegib-Wal comprises homologs of all 15 genes encoding the phage-derived type VI-like secretion system described previously in the amoeba symbiont *Amoebophilus asiaticus* [71, 72]. *Tisiphia* sp. tiOegib-Wal, *Wolbachia* sp. wOegib-WalA, and *Wolbachia* sp. wOegib-WalB encode all components of the conserved Rickettsiales type IV secretion system [53, 73, 74]. In addition, the genome of *Tisiphia* sp. tiOegib-Wal contains the *tra* gene clusters that encode a conjugative DNA-transfer element previously described in several *Rickettsia* genomes [53, 75].

Effectors of intracellular bacteria often harbor domains that have a high sequence similarity with eukaryotic proteins [76]. Proteins including such eukaryotic-like domains target and manipulate different key cellular processes and are used to interact with the host immune response [76, 77]. Further, a correlation between the bacterial lifestyle and the number of eukaryotic-like domains was suggested, where symbionts of microbial eukaryotes, such as amoeba, tend to encode a higher number of proteins with eukaryotic-like domains than free-living bacteria and intracellular bacteria of multicellular hosts [76]. However, the correlation was only systematically analyzed for *Legionella* and Chlamydiae. Here, we extend this analysis to *Cardinium, Wolbachia* and *Rickettsiaceae.* To investigate the number of proteins with eukaryotic-like domains in the symbionts of *O. gibbosus* we screened their proteomes and those of selected relatives for known eukaryotic domains.

The genome of *Cardinium* sp. cOegib-Wal contains only few ankyrin (ANK) and tetratricopeptide repeat (TPR) containing proteins (Table 1). ANK and TPR repeats are part of the so-called tandem-repeat domains that mediate protein-protein and protein-nucleic acid interactions in eukaryotes [76]. The presence of ANK and TPR containing proteins were also described for other group A *Cardinium* strains [49, 78]. *R. oedothoracis* encodes proteins with tandem-repeat domains, proteins with chromatin modulation domains, and a protein with a ubiquitination domain (Table 1). Proteins including chromatin modulation domains are able to influence host gene expression, and ubiquitin is a post-translational modification that mediates various protein interactions in eukaryotes. The genome of *Tisiphia sp. tiOegib-Wal* also contains proteins with all three types of domains (Table 1). The two *Wolbachia* genomes wOegib-WalA and wOegib-WalB contain proteins with domains involved in ubiquitination and a high number of ANK repeat-containing proteins (Table 1). This is in line with the number of ANK repeat-containing proteins found in other arthropod-infecting *Wolbachia* strains [79, 80]. So far no correlation could be found between the number of ANK repeat-containing proteins and the phenotype of *Wolbachia,* i.e. cytoplasmic incompatibility or male-killing [79].

**Table 1:**
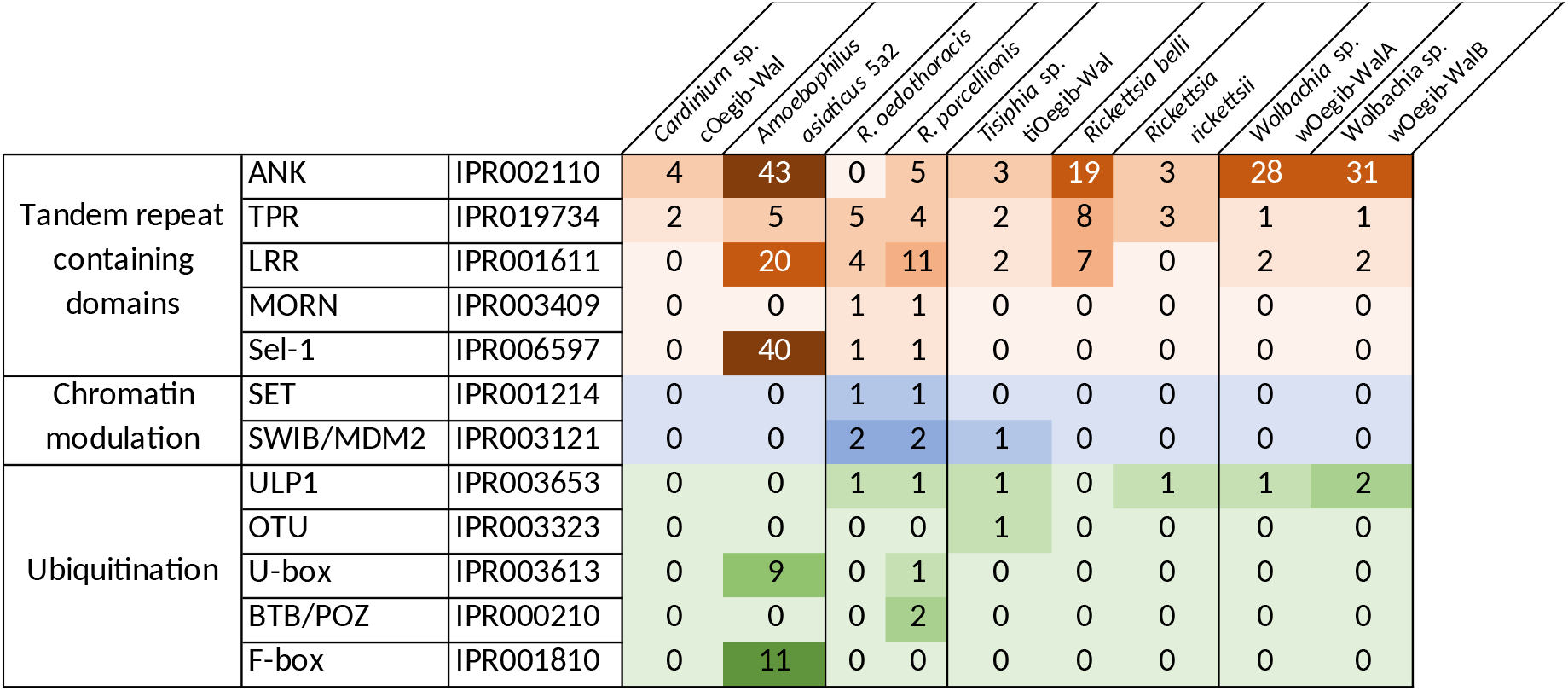
Eukaryotic-like domain containing proteins of the endosymbionts of *O. gibbosus* and selected closely related bacteria.

Extending earlier observations ([76]), our results suggest that amoeba-symbionts like *Amoebophilus asiaticus* (n=128) or *Protochlamydia amoebophila* (n=106) encode a large number of proteins with different eukaryotic-like domains, symbionts that are capable to infect both amoeba and multicellular organisms like *Rickettsia bellii* (n=34), *Waddlia chondrophila* (n=13), or *Simkania negevensis* (n=25) include an intermediate number, and symbionts only found to infect multicellular organism like *Rickettsia rickettsii* (n=7) or members of the family *Chlamydiaceae* (n=6) favor a smaller number of proteins with eukaryotic-like domains (Table 1) [76]. The high number of ANK containing proteins found in *Wolbachia* represents a notable exception, perhaps indicating different selective forces acting on those proteins. Accordingly, the *Rickettsiaceae* and *Cardinium* endosymbionts of *O. gibbosus* would be predicted to be restricted to multicellular hosts, whereas *Rhabdochlamydia* would be able to infect both unicellular and multicellular hosts. This scenario is consistent with members of the genus *Rhabdochlamydia* representing intermediate stages of transition from unicellular to multicellular hosts, as suggested earlier [39].

#### Transposases indicate horizontal transmission and past coexistence of the endosymbionts

Our analysis revealed that the genomes of all five endosymbionts of *O. gibbosus* contain a high number of transposable elements (TEs), especially insertion sequences (ISs). This simplest form of TEs consists only of a transposase gene flanked by inverted repeats [81]. ISs are mobile elements that can spread within a genome independent of cellular replication and are known to be major drivers of bacterial genome evolution and diversification [82]. They are involved in genome size reduction by inactivation of genes under relaxed selection [83], as suggested for *Rhabdochlamydia [39].* ISs can also move between genomes of different species by means of horizontal gene transfer (HGT) and thereby mediate the exchange of genetic material across species [82]. The presence of large numbers of mobile elements is common in genomes of facultative intracellular and extracellular bacteria but was regarded to be limited or absent in ancient obligate intracellular bacteria that thrive in a restricted niche with limited interaction with other bacteria [84, 85]. However, according to the “intracellular arena hypothesis” obligate intracellular bacteria that switch hosts, indeed come into contact with other bacteria and thus can accumulate higher numbers of mobile elements in their genomes than those that are strictly bound to one host [84]. This hypothesis could also explain the high number of TEs found in the spider endosymbionts as even though the genera *Cardinium, Wolbachia,* and *Rickettsiaceae* consist of obligate and mainly maternally transmitted endosymbionts, horizontal transmissions occur in all of them [14, 86–93]. This is also documented by the diverse co-occurrence patterns within and between populations observed here and elsewhere (Figure 1) [21].

As TEs are frequently exchanged between bacteria sharing an environmental niche, we used TE gene trees to infer whether the spider endosymbionts recently shared TEs or have coexisted and shared TEs in the past. For this purpose, we clustered all transposase genes into gene families, extracted those shared by two or more endosymbionts (Table S6), and calculated phylogenies for each of them (Figure 4; Figure S7). We found shared TEs between *Cardinium* sp. cOegib-Wal and *Tisiphia sp. tiOegib-Wal, Tisiphia sp. tiOegib-Wal* and *Wolbachia* wOegib-WalB, and a TE shared between ‘Ca. *Tisiphia’, Cardinium* and *Wolbachia* (Figure 4, Table S6). Sister-clade relationship between the TEs in all phylogenetic trees and the presence of some TEs in other *Cardinium* and *Wolbachia* strains provide evidence for ancient evolutionary links and a possible shared niche in the past. Yet, we did not find any evidence for more recent transfers of transposase genes (Figure 4). This finding is consistent with our results from screening the entire endosymbiont genomes for genes putatively acquired through HGT. While several HGT-derived candidates were predicted by HGTector [94] for all symbionts, we could not find support for any recent HGT events among the spider symbionts. Interestingly, however, TEs of *Wolbachia* wOegib-WalA and *Tisiphia sp. tiOegib-Wal* are related to a TE from the amoeba symbiont *Paracaedibacter symbiosus* (IS110; Figure 4). This suggests an ancient co-existence of these amoeba and arthropod symbionts and is reminiscent of a similar case, in which a TE (ISRpe1) was found to be shared by *Wolbachia, Cardinium*, ‘Ca. *Tisiphia*’, and the amoeba symbiont *Amoebophilus asiaticus* [95]. Taken together, our phylogenetic analysis of transposase genes provides evidence for a shared niche of the *O. gibbosus* symbionts with amoeba symbionts and a co-occurrence during their evolutionary history. This analysis also shows that, due to their frequent transmission across species borders, TEs may provide a promising tool to reconstruct historical associations of microbes.

**Figure 4:**
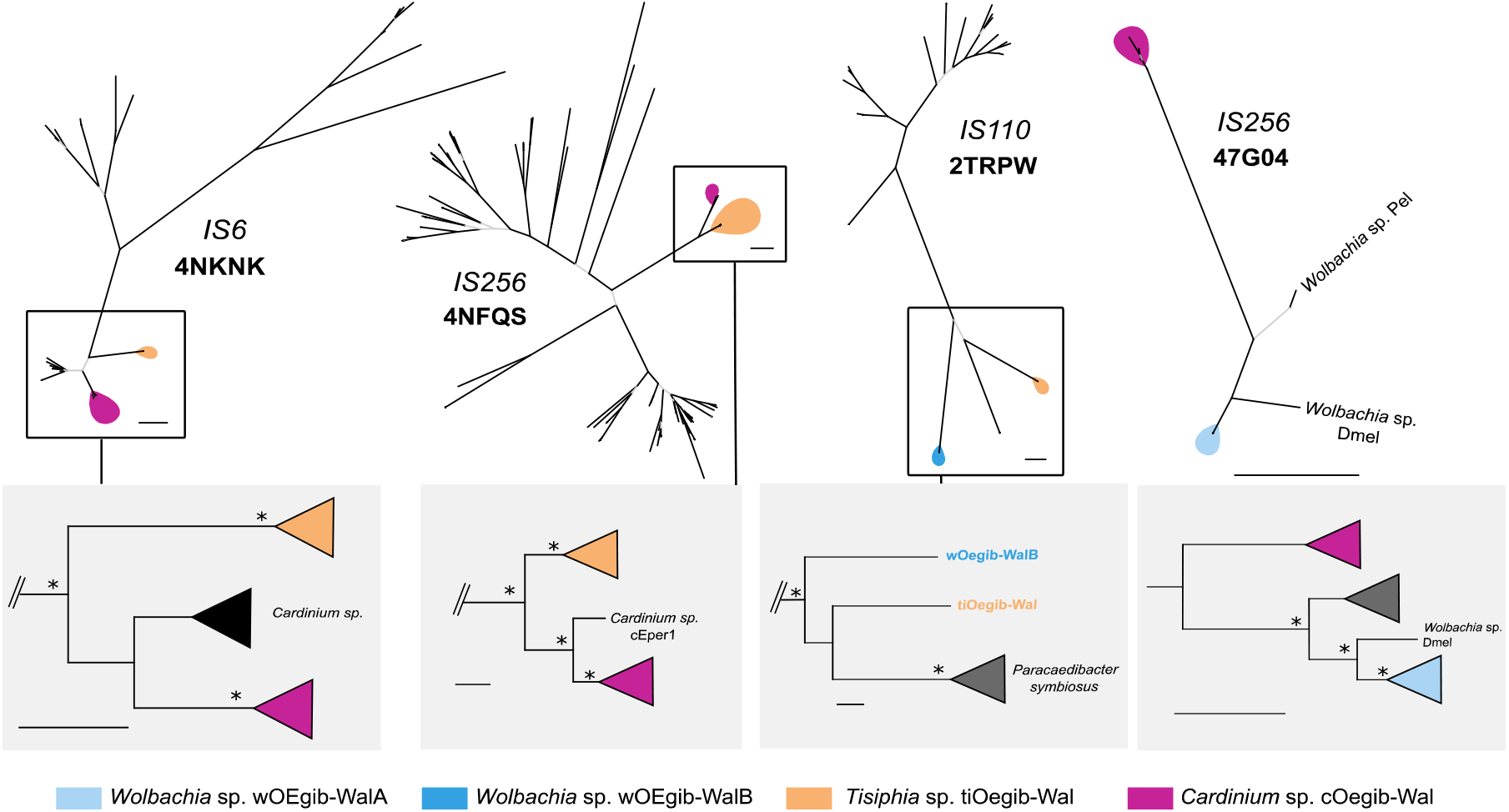
Unrooted phylogenies of shared transposases of the endosymbionts. Branches with bootstrap values >95 are indicated in black, branches with lower bootstrap values are shown in light gray. The trees are labeled with the respective IS-family and eggNOG OG. Below each phylogeny, midpoint rooted trees of selected clades are shown. Branches with support values >95 are indicated with asterisks. Further details on the phylogenies can be found in table S6. Note that for the trees only complete sequences were used, thus in some cases not all endosymbionts stated in table S6 are represented in the trees. Scale bar indicates 0.1 substitutions per position in the alignment.

## Conclusions

In the current study we could show that *O. gibbosus* is co-infected with up to five different endosymbionts that are heterogeneously distributed among different host populations. In contrast to other studies on arthropod endosymbionts that suggested co-infections with more than two endosymbionts occurred only rarely, we found that about 30% of the 16 *O. gibbosus* individuals from six populations analyzed here were infected with three or more endosymbionts. We were able to reconstruct four complete endosymbiont genomes present in the same host population. This offered us the unique opportunity to holistically explore this host-endosymbiont system on a genomic level. We identified all endosymbionts as facultative symbionts that are not essential for the survival of the spider and found indications for horizontal transmission of the endosymbionts, a route that is proposed for facultative symbionts as an alternative to the primarily vertical transmission [14, 86–93]. We detected large numbers of transposable elements in all endosymbiont genomes and could show that ancestral *Cardinium*, ‘*Ca.* Tisiphia’ and *Wolbachia* endosymbionts may have coinfected the same hosts in the past by using phylogenetic reconstructions of shared transposable elements. In summary, our findings contribute to broadening our knowledge about important groups of endosymbionts and set the stage for more in-depth analysis of host-endosymbiont interactions and interactions between different endosymbionts using the spider *O. gibbosus* as a model system.

## Methods

### Endosymbiont genome assembly

To reconstruct the endosymbiont genomes, we used long-read (Pacbio; W744×W766) and short-read (Illumina; W791) sequencing libraries from the host *O. gibbosus* described in [96]. As references for the long-read alignments, we used metagenome-assembled genomes (MAGs) from the endosymbionts reconstructed from the short-read libraries and contigs from the host-assembly classified as endosymbiont contigs using a custom Kraken database as described in [96].

For *Cardinium* sp. cOegibbosus-W744×776, *Wolbachia* sp. wOegibbosus-W744×776A and *Wolbachia* sp. wOegibbosus-W744×776B we mapped long reads to the respective MAGs and contigs using minimap2 (v2.17) [97]. Finally, all mapped reads were merged, and duplicates were removed. As the coverage was too high for *Wolbachia*, the mapped reads were subsampled to a coverage of 70x. The final set of reads was then assembled using unicycler (v0.4.6) [98]. The quality of the assemblies was checked by checkM (v1.0.18) [99] and visually inspecting the assembly graph [100]. The *Wolbachia* genomes were closed manually, and the assembly of *Wolbachia* sp. wOegibbosus-W744×776B was manually curated to get rid of misassembled regions.

For *Candidatus* Tisiphia sp. Oegibbosus-W744×776 we mapped long reads to the respective MAG and contigs using minimap2 (v2.17) [97] and short reads using bbmap (v37.61) (sourceforge.net/projects/bbmap/). Both long and short reads were merged and deduplicated afterwards. The final set of long and short reads was then used for a hybrid assembly using unicycler (v0.4.6) [98]. The quality of the assembly was checked by checkM (v1.0.18) [99] and visually inspecting the assembly graph [100].

*Rhabdochlamydia oedothoracis* MAG OV001 was assembled using the short-read library from the Overmere_OV001 dataset. We first carried out a *de-novo* assembly using megahit (v1.1.2) [101]. Afterwards, the contigs were binned using metabat2 (v2.15) [102]. The required bam file for binning was created using bowtie2 (v2.3.5.1) [103]. Finally, the qualities of all bins were checked with checkM (v1.0.18) [99]. The bin belonging to *Rhabdochlamydia* was used for a reassembly. For that purpose, we mapped the short-read library from Overmere_OV001 to the *Rhabdochlamydia* bin and the reference genome *R. oedothoracis* W744×W776 (GenBank accession number: CP075587-CP075588) using bbmap (v37.61) (sourceforge.net/projects/bbmap/). Afterwards all mapped reads were merged, and duplicates removed. This set of reads was then used for a second round of *de-novo* assembly and binning as described before. The quality of the final MAG was again checked with (v1.0.18) [99].

Origins of replication were detected using originx [104] and Ori-Finder [105]. The orientation of genome sequences was adjusted to previously sequenced *Wolbachia* and *Rickettsia* genomes [106].

### Abundance estimations of the endosymbionts

To estimate the abundance of the different endosymbionts in *O. gibbosus* individuals we used a dataset consisting of 16 short-read (Illumina) sequencing libraries obtained from male *O. gibbosus* individuals from six different sampling locations (Damvallei, n=4; Sevendonck, n=2; Pollismolen, n=2; Honegem, n=2; Overmeren, n=2; Walenbos, n=4) described in [96]. The sequencing libraries are deposited in NCBI (BioSample: SAMN16961399 - SAMN16961414). Illumina reads were right-tail clipped using a minimum quality of 20, and all reads shorter than 75 after this treatment were discarded using FASTX-Toolkit (http://hannonlab.cshl.edu/fastx_toolkit/). Reads with undefined nucleotides or left without a pair were removed using PRINSEQ [107]. Afterwards, the reads were mapped to the genome of *O. gibbosus* (GCA_019343175.1) using bbmap (v37.61; “-minid=99”) (sourceforge.net/projects/bbmap/). The unmapped reads were then mapped to the endosymbiont genomes, and the mapped reads to elongation-factor alpha (EF1alpha) from *O. gibbosus* using bowtie2 (v2.3.5.1) [103]. The median coverage for each endosymbiont and the mean coverage for EF1alpha per sample was calculated using samtools (v1.12) [108]. For ‘*Ca.* Tisiphia’, *Cardinium,* and *Rhabdochlamydia* the coverage across the whole genome was used, for *Wolbachia* A and *Wolbachia* B only the coverage of the respective accessory genes. The median coverage of the endosymbionts was then normalized by the mean coverage of EF1alpha per sample.

### Phylogenetic analyses

For the phylogenetic reconstructions of *Wolbachia* and *Cardinium* we downloaded representative genomes belonging to different groups from NCBI (Table S7). In the case of *Wolbachia, Anaplasma phagocytophilum* strain HZ and *Ehrlichia canis* strain Jake were used as outgroups and for *Cardinium, Amoebophilus asiaticus* strain 5a2 was used. Ribosomal RNA proteins were extracted and universally conserved ones selected. The protein sequences were individually aligned using MAFFT (v7.453; “--localpair” “--maxiterate 1000”) [109]. Divergent and ambiguously aligned blocks were removed using Gblocks (v0.91b) [110]. The alignments were then concatenated using custom perl scripts and Bayesian inference was performed using MrBayes (v3.2.7) [111], using the JTT+I+G4 substitution model. We ran two independent analyses with four chains each for 300,000 generations and checked for convergence (convergence diagnostic <=0.01).

For *Rickettsiaceae* we also downloaded representative genomes from NCBI and used *Megaira* sp. strain MegNEIS296, *Megaira* sp. strain MegCarteria, *Occidentia massiliensis* strain Os18, *Orientia tsutsugamushi* strain Boryong, and *Orientia chuto* strain Dubai as outgroups (Table S7). We extracted ribosomal RNA proteins and selected universally conserved ones. The protein sequences were individually aligned using MUSCLE (v3.8.31) [112] and the alignments were concatenated using AMAS [113]. The phylogeny was calculated with iqtree2 (v2.1.2; “-bnni” “-alrt 1000” “-m TESTNEW” “--madd LG4X” “-bb 1000”) using the JTTDCMut+F+R3 substitution model [114]. In order to verify if the endosymbionts belong to an existing clade or represent a new one we calculated the average amino acid and average nucleotide identity for the endosymbiont and selected representative genomes using the method described by [41].

### Genome annotation and genome content analysis

The genome of *‘Candidatus* Tisiphia’ sp. Oegibbosus-W744×776, *Rhabdochlamydia oedothoracis* MAG OV001, *Wolbachia* sp. wOegibbosus-W744×776A and *Wolbachia* sp. wOegibbosus-W744×776B were annotated using prokka (v 1.14.6; “--compliant” “--kingdom Bacteria”) [115]. The genome of *Cardinium* sp. cOegibbosus-W744×776 was annotated using Prokka (v1.14.6; --kingdom Bacteria) [115], infernal (v1.1.4) with the bacterial subset of Rfam (v14.2; --cut_tc -mid) [116], and tRNAscan-SE (v2.0.9; -B -I -isospecific) [117]. The NRPS gene was identified from the draft prokka annotation, and a more detailed annotation was done using AntiSMASH (v6.1) as implemented through the webserver [118].

For the genome content analysis of ‘*Ca.* Tisiphia’, *Cardinium* and *Wolbachia,* we mapped all proteins against the eggNOG database (v5.0) [119] using emapper (v2.1.0) [120] to assign them to existing gene families. For *Rhabdochlamydia,* we used the eggNOG database (v4.5.1) [42] and emapper (v1.0.1, “-d bact”) [121]. For all unmapped proteins we performed an all-against-all blastp [51] search and clustered proteins with an e-value < 0.001 *de-novo* with SiLiX (v1.2.11) [43] with default parameters. For functional annotation we used eggNOG (v5.0; v4.5.1) [42, 119], and blastp against the NCBI nr database for the *de-novo* gene families. For the analysis of the accessory genomes we removed all proteins shorter than 100 amino acids in order to remove ghost CDSs and remnants of transposases. To show the co-localization of transposase genes, accessory genes and regions covered by *R. oedothoracis* MAG OV001, we extracted the loci of transposase and accessory genes using a custom python script and identified homologous regions between *R. oedothoracis* W744×776 and MAG OV001 using mummer (v3.0; “nucmer --mum”; “delta-filter -qr”) [122]. Afterwards, the genome was plotted with circos [123]. To link breaks in the synteny between *R. oedothoracis* W744×776 and *R. oedothoracis* MAG OV001 to transposases, contigs showing rearrangements were selected using mauve. Homologies between the contigs and the reference genome were identified using blastn and the alignments and annotations were visualized using R (v4.0.3) [124] and the genoPlotR package (v0.8.11) [125].

Secretion systems were identified by blastp (“-evalue 1e-3”) [51] against selected representatives *(Cardinium* sp. cEper1 (GCA_000304455.1), *Tisiphia* sp. RiCNE (GCA_002259525.1), *Wolbachia* sp. wMel (GCA_000008025.1)). The protein sequences were then individually aligned using MAFFT (v7.453; “--maxiterate 1000”) (Katoh & Standley, 2013). To investigate the presence of eukaryotic-like domains in the endosymbiont genomes, we screened their proteomes and those of selected representatives *(Amoebophilus asiaticus* 5a2 (GCA_000020565.1), *Rhabdochlamydia porcellionis* (GCA_015356815.2), *Rickettsia bellii* RML369-C (GCA_000012385.1), *Rickettsia rickettsii* Iowa (GCA_000017445.3)) using InterProScan (v5.53-87.0; “-appl Pfam” “-dp”) (Jones *et al.,* 2014). The presence of vitamin and essential amino acid biosynthesis pathways was checked using GhostKOALA (V2.2) [126] and the “reconstruct” function of KEGG Mapper [127]. Biotin genes in the *Wolbachia* genomes were identified as described for the secretion systems *(Wolbachia* sp. wNfla (GCA_001675695.1), *Wolbachia* sp. wNleu (GCA_001675715.1), *Wolbachia* sp. wLug (GCA_007115045.1), *Wolbachia* sp. wCle (GCA_000829315.1), *Wolbachia* sp. wstri (GCA_007115015.1)). Toxin-antitoxin systems were identified by text search (grep -i “toxin\ļantidote”) against the emapper and prokka annotations. The results were checked by blastp against the NCBI nr database.

### Analysis of shared transposases

For the phylogenetic analysis of shared transposases we first clustered all genes annotated as transposases by prokka into gene families using SiLiX (v1.2.11) [43]. For each gene family that was shared by two or more endosymbionts we searched for homologous sequences using the blastp function of ISfinder [128] and created a multiple sequence alignment with MAFFT (v7.453; “--maxiterate 1000”). Afterwards, the alignments were manually checked and sequences showing clear signs of degradation either on the 3’ or 5’ end were removed. Finally, the alignments were trimmed using bmge (v1.12) [129] and used for phylogenetic reconstruction using iqtree2 (v2.1.2; “-bnni” “-alrt 1000” “-m TESTNEW” “-bb 1000” “-mset LG” “-madd LG+C10,LG+C20,LG+C30,LG+C40,LG+C50,LG+C60” “-keep-ident” “-wbtl”) [114]. For transposase sequences showing a clear sister-clade relationship in the *de-novo* trees and belonging to the same eggNOG gene family, we calculated phylogenetic trees using gene families based on the eggNOG database (v5.0) [119] and emapper (v2.1.0) [120]. For this purpose, we added the protein sequences from the respective gene families from the eggNOG database (v5.0)) [119] to the endosymbiont gene families. For each gene family we then calculated multiple sequence alignment, curated them and reconstructed phylogenies as described above.

### Prediction of HGTs

We used HGTector (v2.0; “--evalue 1e-20 --identity 30 --coverage 40”; and for *‘Ca. Tisiphia’* “-self-tax 114295 --close-tax 780”) [94] to predict horizontal gene transfer events in the endosymbiont genomes. To keep the analysis conservative, we excluded all genes annotated as transposable elements (IS elements, phages, introns) or genes that are part of TEs (integrases, reverse transcriptases) from the output. We also excluded genes predicted to be transferred from members of the phylum Chlamydiae for *Rhabdochlamydia oedothoracis* W744×776 and genes predicted to be transferred from members of the family *Rickettsiaceae* for *Candidatus* Tisiphia sp. Oegibbosus-W744×776 as we focused our analysis on HGTs between distantly related groups.

### SNP calling

For calling genetic variants we used the same short-read sequencing libraries as described for the abundance estimations, but we excluded all samples from the Walenbos population as the endosymbiont genomes were reconstructed from this population, and we excluded all samples with a coverage < 30. Afterwards, we applied breseq (v0.36.1) [130] to call genetic variants present in the sequencing libraries in comparison to the reference genomes. We excluded all variants with a frequency < 1 and variants affecting transposases or introns from the output and calculated the total number of mutations/kB and relative abundances of variant types per sample.

### Statistical analysis

All statistical tests and data analysis were performed in R (v4.0.3) [124] and visualized using ggplot2 (v3.3.3) [131]. The hierarchical clustering was done using the “hclust” function (“stats’’ package v4.0.3) and a distance matrix created from symbiont abundance data using the “dist” function (“stats’’ package v4.0.3).

## Acknowledgements

The authors would like to acknowledge the staff maintaining the *Life Science Compute Cluster* (LiSC; http://cube.univie.ac.at/lisc) at the University of Vienna which was used for computational analysis.

## Author contributions

T.H., S.K., M.H. conceptualized this study. T.H. and A.M.M. performed phylogenetic and comparative genomics analyses. T.H. performed abundance and SNP analysis and phylogenetics of transposases. T.H., A.M.M., S.K. and M.H. interpreted the results. F.H., S.K., A.M.M. and T.H. performed sequencing and assembly of the endosymbiont genomes. F.H. performed sequencing of the *O. gibbosus* individuals from different populations. All authors wrote and edited the paper.

