## Supplementary Methods and Figures S1-S7 for "One to host them all: genomics of the diverse bacterial endosymbionts of the spider *Oedothorax gibbosus*"

To calculate the 16S rRNA gene phylogeny for the genus '*Candidatus* Tisiphia' we downloaded all available *Rickettsia* and '*Candidatus* Tisiphia' 16S rRNA gene sequences from SILVA SSU (r138.1) on 16 November 2021. Further, we added 16S rRNA gene sequences from the dataset we used for the ribosomal protein gene phylogeny (Table S7) and sequences obtained from other arachnid hosts [1]. We then removed duplicates and filtered the sequences to remove sequences shorter than 1,000 bp. The remaining sequences were aligned using SINA ("variability profile: bacteria") [2] and trimmed using trimAl (v1.4.rev15; "-noallgaps") [3]. The phylogeny was then calculated with iqtree2 (v2.1.2; "-bnni" "-alrt 1000" "-m TESTNEW" "--madd LG4X" "-bb 1000") using the TVMe+R6 substitution model [4].

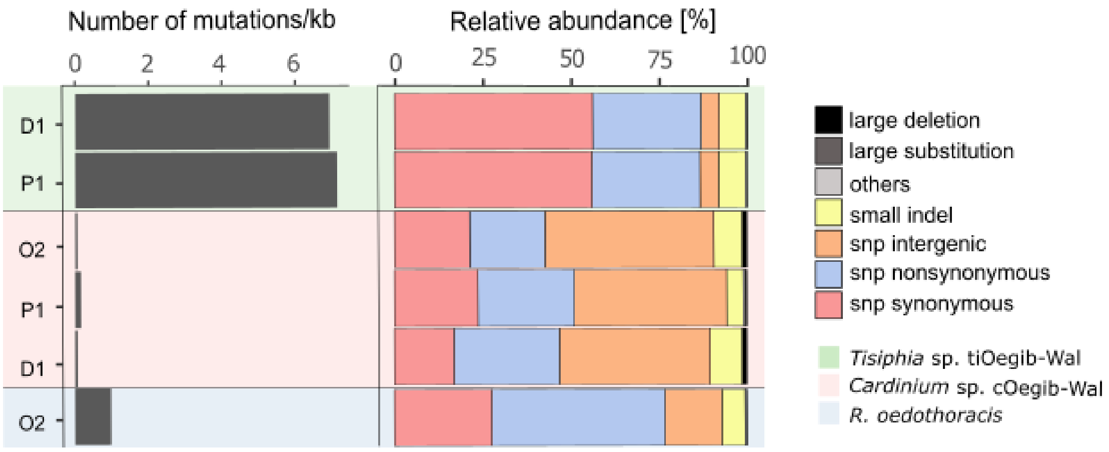


Supplementary figure S1: Frequency and relative abundance of genomic variations found in '*Candidatus* Tisiphia', *Cardinium* and *Rhabdochlamydia* endosymbionts of selected *O. gibbosus* populations.

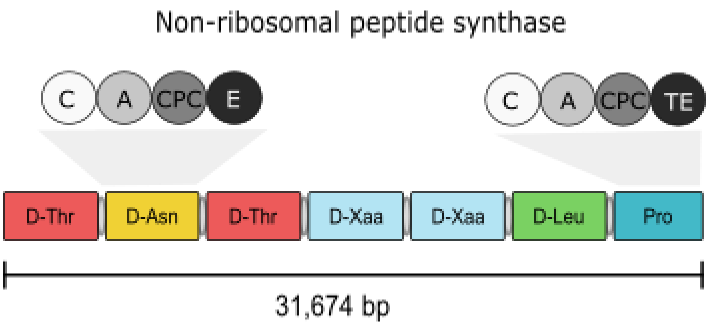


Supplementary figure S2: Structure of the non-ribosomal peptide synthase in the genome of *Cardinium* sp. cOegib-Wal. The NRPS consists of seven modules, each incorporating one amino acid (Thr - Threonine, Asn - Asparagine, Leu - Leucine, Pro - Proline, Xaa - Any amino acid). Each module consists of four domains: an adenylation domain (A), an peptidyl carrier protein domain (PCP), a condensation domain (C) and an epimerization domain (E). The last module contains a thioesterase domain (TE) instead of an epimerization domain.

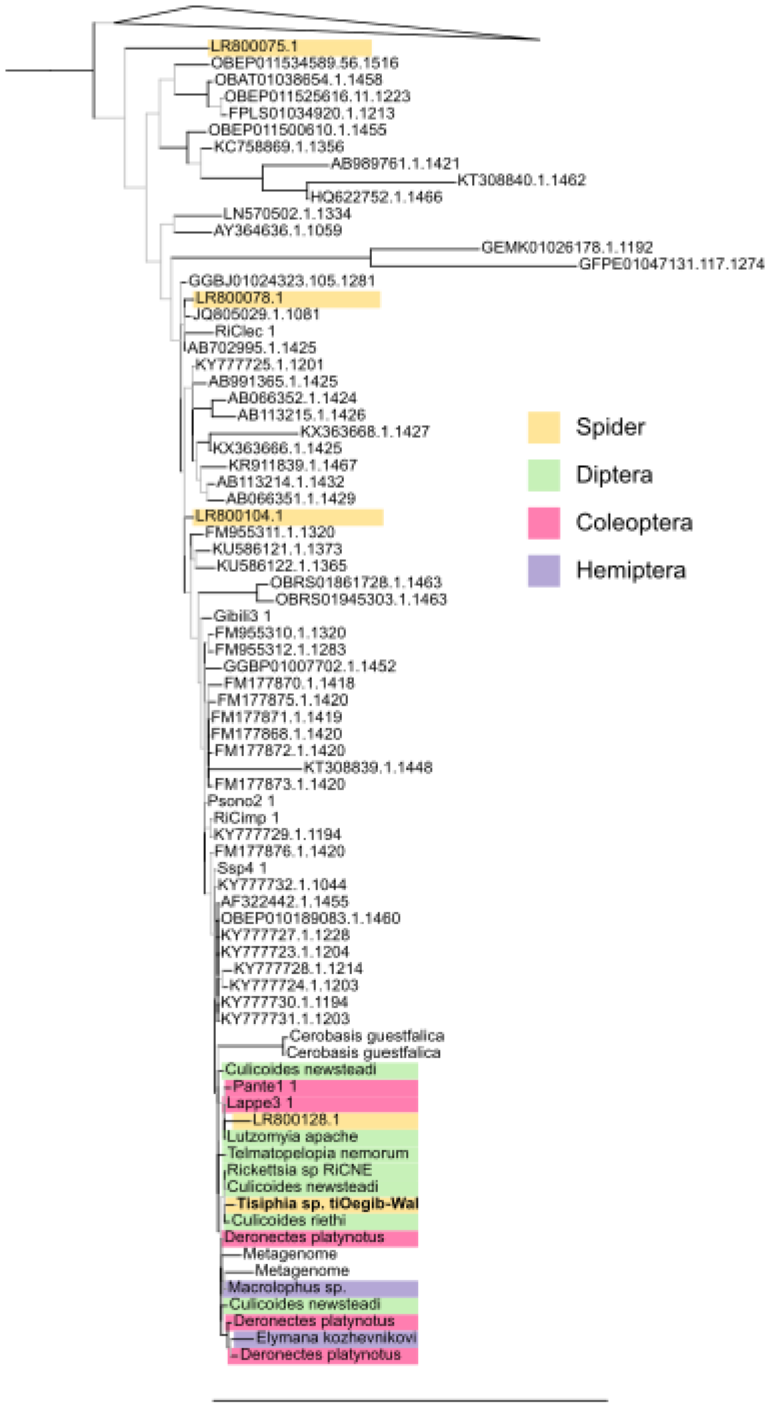


<u>Supplementary figure S3</u>: 16S rRNA gene phylogeny of the genus '*Candidatus* Tisiphia'. Scale bar indicates 0.1 substitutions per position in the alignment. The tree was rooted using *Orientia* sp. and '*Candidatus* Megaira' sp. as an outgroup for visualization. Branches with bootstrap values >95 are indicated in black, branches with lower bootstrap values are shown in light-gray. For visualization sequences belonging to the genus *Rickettsia* were collapsed.

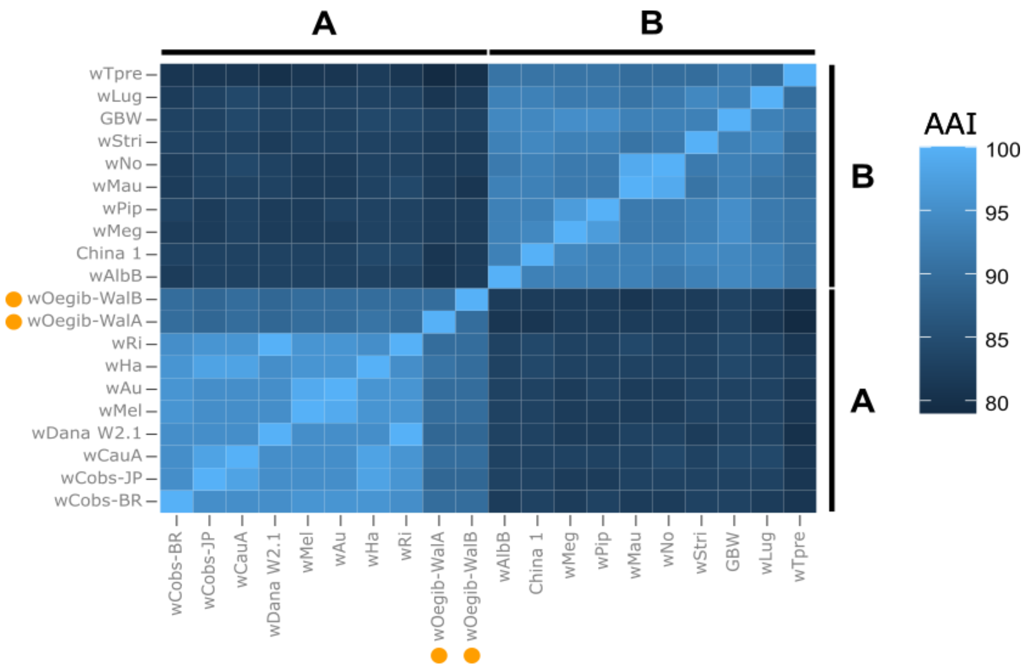


Supplementary figure S4: Average Amino Acid Identity (AAI) of different members of supergroup A and B of the genus *Wolbachia*. The AAI was calculated as described in [5].

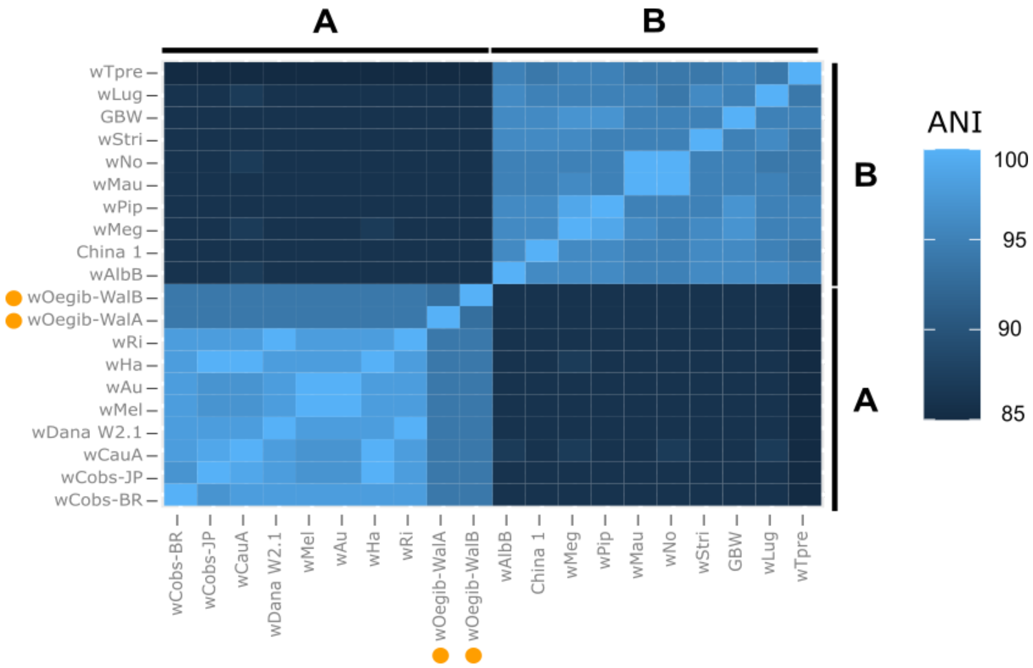


Supplementary figure S5: Average Nucleotide Identity (ANI) of different members of supergroup A and B of the genus *Wolbachia*. The ANI was calculated as described in [5].

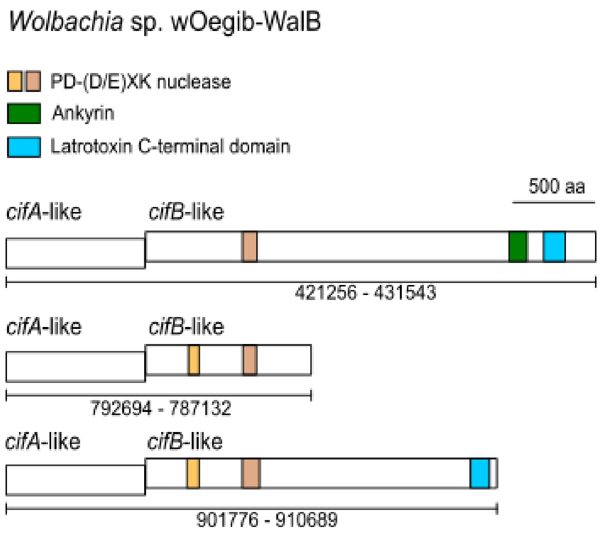


Supplementary figure S6: Analysis of putative cif gene clusters in *Wolbachia* sp. wOeGib-WalB. Remote homologs of cifA- and cifB-like genes potentially involved in cytoplasmic incompatibility in the genome of wOegib-WalB and identification of typical domains.

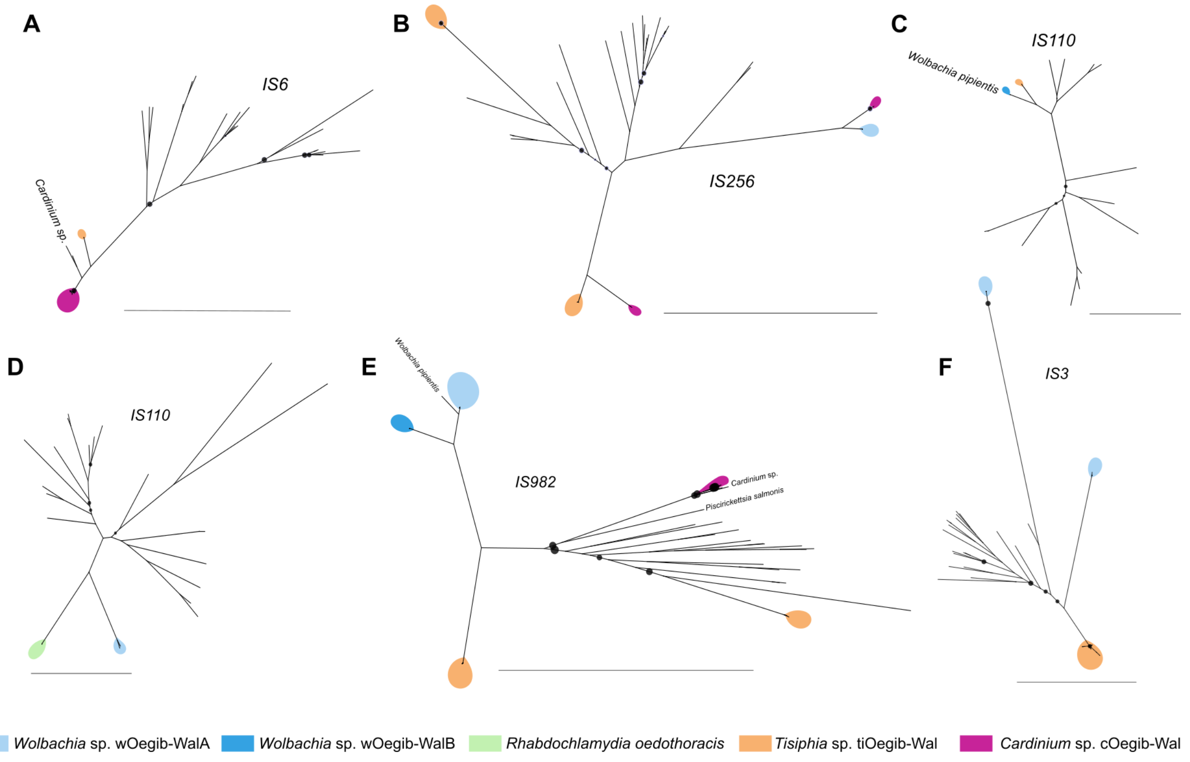


Supplementary figure S7: Phylogenies of *de-novo* clustered TE gene families. Scale bar indicates 0.1 substitutions per position in the alignment. Black dots indicate bootstrap values, where the size indicates values ranging from 0-95. Further details on the gene families can be found in Table S6. Note that for the trees only complete sequences were used, thus in some cases not all endosymbionts stated in Table S5 are represented in the trees.
